# Dynamic Control of Eye-Head Gaze Shifts by a Spiking Neural Network Model of the Superior Colliculus

**DOI:** 10.1101/2022.08.10.503437

**Authors:** Arezoo Alizadeh, A. John Van Opstal

## Abstract

To reorient gaze (the eye in space) towards a target can be achieved by infinitely many combinations of eye- and head movements. However, behavioral measurements show that the primate gaze-control system selects specific contributions of eye- and head to the saccade, which depend on the initial eye-in-head orientation. Single-unit recordings in the primate superior colliculus (SC) during head-unrestrained gaze shifts have further suggested that cells may encode the instantaneous trajectory of a desired straight gaze path in a feedforward way by the total cumulative number of spikes in the neural population, and that the instantaneous gaze kinematics are determined by the neural firing rates. The recordings also indicated that the latter depended strongly on the initial eye position as well. We recently proposed a conceptual model that accounts for many of the observed properties of eye-head gaze shifts and on the potential role of the SC. Here, we extend and test the model by incorporating a spiking neural network of the SC motor map, the output of which drives the eye-head motor control circuitry by linear cumulative summation of individual spike effects of each recruited SC neuron. We propose a simple neural mechanism on SC cells that explains the modulatory influence of feedback from an initial eye-in-head position signal on their spiking activity. The same signal also determines the onset delay of the head movement with respect to the eye. The downstream eye- and head burst generators were taken to be linear, as our earlier work had suggested that much of the nonlinear kinematics of saccadic eye movements may be due to neural encoding at the collicular level, rather than at the brainstem. We show that the spiking activity of the SC population drives gaze to the intended target location within a dynamic local gaze-velocity feedback circuit that yields realistic eye- and head-movement kinematics and dynamic SC gaze-movement fields.

## 1. Introduction

### Background

A saccadic gaze shift is the rapid re-orienting movement of the eyes and head that brings the image of a peripheral visual stimulus of interest onto the fovea. Because of the different plant dynamics of the two motor systems, and the limited oculomotor range, not all eye-head combinations are possible or equally efficient in reorienting gaze. Typically, the amplitudes of eye- and head movements are coupled: small gaze shifts are associated with small head movements, and large gaze shifts with larger head movements^[1–3]^. However, the relative contributions of eyes and head to the gaze shift also depend strongly on the initial eye orientation: if the head is initially oriented straight ahead, and the eye-in-head looks contralaterally from the target, the gaze shift will consist of a larger eye- (and hence smaller head-) movement, than when the eye looks in the ipsilateral target direction. Because the eye saccade is much faster than the head saccade, the former gaze shift will be much faster than the latter, as the eye- and head movements are not executed independently, but interact^[1–4]^.

### Kinematics

The differences in head-movement amplitude and overall gaze kinematics relate strongly to differences in the head-movement onset delay with respect to the eye movement. In the contralateral condition, the head starts later than in the ipsilateral situation and can therefore only briefly (if at all) interact with the ongoing eye movement. However, when eye and head start nearly simultaneously, the interaction will be complete, and the slower head movement will reduce the eye velocity, and hence the overall gaze velocity. As a result, the well-known stereotyped main-sequence relationship of eye-only saccades^[5]^ is no longer valid for combined eye-head gaze shifts: when a head movement accompanies the gaze shift, the gaze-peak velocity will be reduced, even though the gaze amplitude may remain the same^[1,2]^.

### Superior Colliculus

Single-unit recordings in the midbrain Superior Colliculus (SC) of head-restrained^[6,7]^ as well as head-unrestrained^[8]^ monkeys have suggested that the population of saccade-related cells encodes a desired straight gaze trajectory by the instantaneous cumulative spike count, whereby the gaze kinematics are determined by the instantaneous neural firing rates.

To illustrate this, Fig. 1 exemplifies the results from a single-unit SC recording from a monkey making large head-unrestrained gaze shifts in and near the cell’s response field. Gaze saccades started from the central fixation point at straight ahead with the eye-in-head in one of three possible initial orientations: 18 deg left (contralateral), straight ahead, or 18 deg right (ipsilateral)^[8]^. Visual targets appeared (mostly) in the right hemifield, in and around the cell’s response field. The top row (Fig. 1A-C) provides the raw data of 371 trials. Panel 1A shows the spiking patterns for each gaze shift, aligned with gaze-saccade onset (yellow line at *t*=0). The cell rapidly increases its firing rate around 20 ms before saccade onset (red line; the cell’s lead time). In Fig. 1B, the number of spikes emitted during each gaze shift has been color-coded, clearly showing the restricted set of gaze vectors to which the cell is tuned. The center of the movement field (the hot spot) is estimated at around [R,Φ] = [37, 20] deg (yellow dots). In Fig. 1C, the instantaneous firing rates (red) and gaze track velocities (black) are shown for 25 individual trials. Note that for many of the trials there is a strong correlation between the two profiles, even though neural firing rates can be quite noisy. In our earlier work we argued (and showed) that this property betrays tight synchronization of SC bursts in the population during saccades^[7,13]^. For this recording, the correlations were *r* > 0.7 for 219/371 trials.

**Figure 1.**
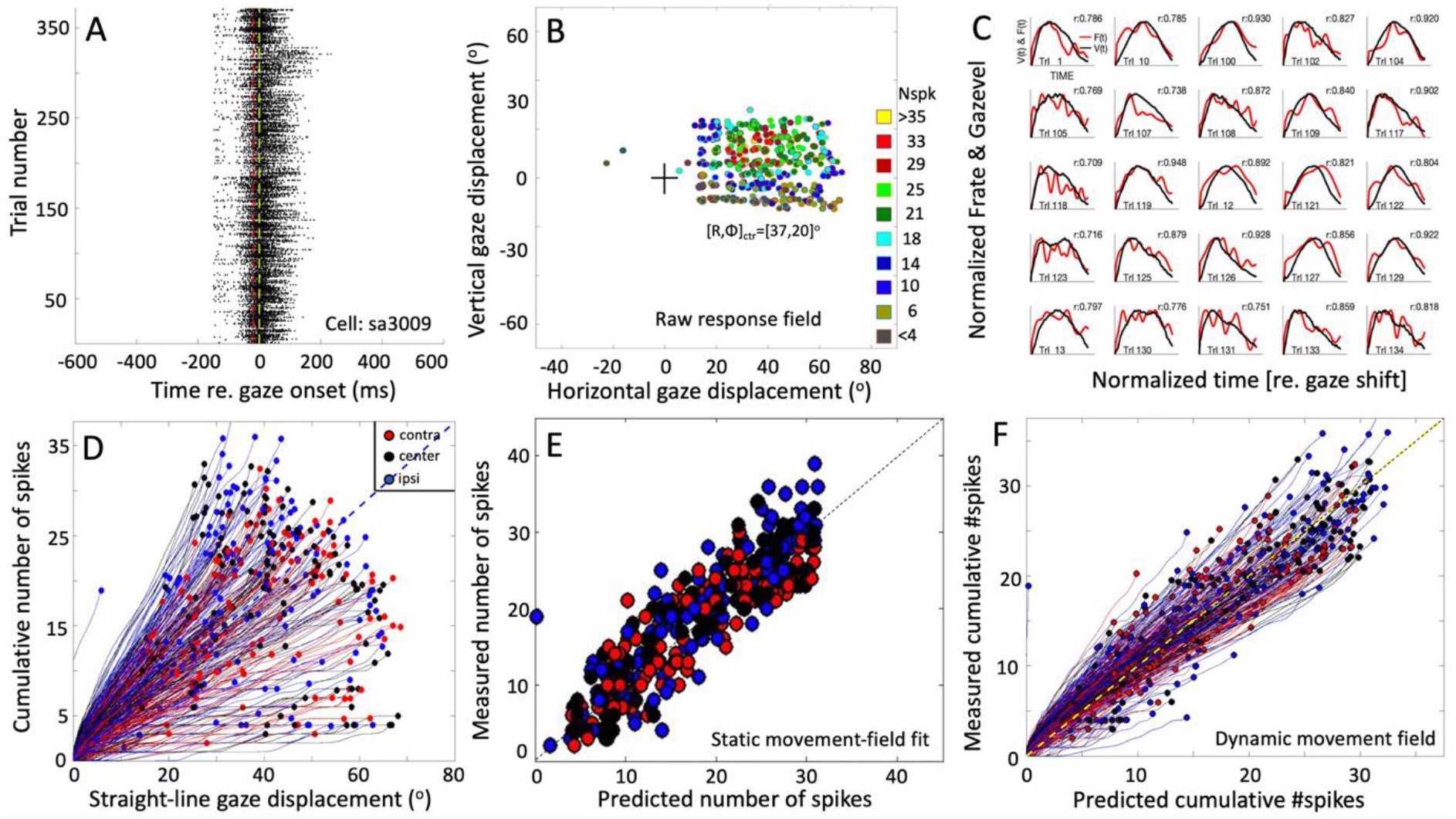
Response properties of SC neuron sa3009 for eye-head gaze shifts into and near its movement field. (A) Dot display of the SC bursts (371 trials) aligned with gaze-shift onset (yellow line at 0). (B) Raw response field of the cell indicates that it is recruited for large rightward gaze shifts (between 20-70 deg) with a small upward component. (C) There is a high correlation between instantaneous firing rate and gaze velocity (shifted backward by 20 ms; shown for 25 trials). For visual purposes, time axes were normalized to saccade durations and amplitudes normalized to their maximum. (D) The straight instantaneous gaze trajectory is linearly related to the cumulative number of spikes in the burst (thin lines). This holds for all gaze shifts (fast and slow). The slopes of the lines vary systematically with the gaze-shift vector. Colors denote the three different initial eye positions in each trial (legend). Note that ipsilateral trials (blue) tend to elicite slightly more spikes than contralateral trials (red). (E) Result of fitting Eqn. 6 (Methods) to the number of spikes in the burst. Parameters: N_0_=31 spikes, R_opt_=36.6°, Φ_opt_=22.3°, σ_p_=0.63 mm, and ε=0.0024 spikes/deg. Correlation fit vs. data: r=0.92. The small eye-position sensitivity, ε, accounts for the eye-position effect seen in (D). (F) Result of Eqn. 8 (Methods) for the dynamic data (r=0.95 for N=37,679 data points).

The cell’s movement field can be modeled quantitatively by relating the total number of spikes in the burst to the gaze vector, incorporating the well-established idea that the gaze shift is encoded by a localized (Gaussian) population within the SC motor map^[9]^. Our group has extended this so-called static ensemble-coding model^[9,10]^ by also accounting for the instantaneous behavior of the firing rates^[6–8]^, linking the cumulative number of spikes of a cell during the gaze shift to the intended straight-line displacement of the eye along the trajectory (delayed by the 20 ms neural lead time). The results of this analysis are shown in Fig. 1D-F (details are described in Methods).

### Models

To understand how a population of SC neurons, like the one Fig. 1, could determine the rich repertoire of eye-head gaze trajectories and kinematics, we recently proposed a gaze-control model in which the SC population specifies the desired straight gaze trajectory by its total cumulative spike count, and saccade kinematics by the firing rates to rapidly drive eye and head to the target within a common dynamic gaze-feedback loop^[8,11]^. This local feedback circuit embodies the interaction between the ongoing eye- and head-motor commands, and hence the strong influence of both the eye- and head kinematics on each other and on the current gaze velocity. The model included the observed influence of initial eye position on the head contribution to the gaze shift and on the output of the SC.

### This study

However, in the model, the SC burst was lumped into a simplified rectangular pulse, rather than by a more realistic distributed code of spike trains from a localized, tuned neural population in the motor map. To explain how the SC neuronal population could control gaze shifts under a variety of initial conditions, we here extended and tested the model by incorporating a two-layer spiking neural network, in which the neurons displayed similar firing behaviors as has been recorded in monkey (Fig. 1) ^[7,8]^. The SC activity is thought to issue a feedforward desired straight eye-head gaze trajectory through dynamic linear summation of the individual spike effects to a downstream gaze-feedback comparator^[6–8]^. To account for the influence of initial eye-in-head orientation on the observed firing characteristics of SC neurons, we here propose a single neural mechanism by which a feedback oculomotor signal modulates two sensitivity parameters of all neurons in the motor map to affect their bursting behavior.

Note that because of several nonlinearities in the brainstem circuitry that control the eye- and head-movement kinematics, like a limited oculomotor range, activation of the vestibulo-ocular reflex, and the varying delay of head-movement onset and resulting head-movement contributions to the gaze shift^[1–3]^, the simple linear relationship between SC firing rates and movement kinematics, as shown for head-restrained ocular saccades^[6]^, might be expected to break down. However, the example of Fig. 1D seems to suggest that this relationship may still hold. To better understand these properties, we also analyzed the results in detail by determining the dynamic SC movement fields of our model neurons for simulated gaze shifts of different amplitudes from different initial eye orientations^[6,8]^, and compared the data with single-unit recordings from monkey SC neurons (Fig. 1) and with actual monkey gaze shifts.

## 2. Methods

### Electrophysiological recordings

The monkey experiments (Figs. 1 and 9B) were performed in the laboratory of dr. EG Freedman at the Department of Neurobiology and Anatomy, School of Medicine and Dentistry of the University of Rochester, NY, while one of the authors (AJVO) was a visiting scientist. Two rhesus monkeys (P and S) took part in these experiments. They had been trained to follow a briefly flashed visual target with a fast eye-head gaze shift, while single-unit activity was recorded from the left SC (rightward gaze saccades). Animals received a small liquid reward for each successful trial. Details on surgical procedures, training protocols, and experimental setup are described in full detail elsewhere^[12–14]^. Additional details on the experimental paradigm are provided in the Supplemental Material. Experimental procedures and protocols were all approved by the University of Rochester Animal Care and Use Committee, and fully adhered to the National Institutes of Health Guide for the Care and Use of Animals. We recorded from a total of 52 cells, out of which 30 neurons could be isolated sufficiently long for a detailed analysis. The movement fields were typically obtained from cells in the caudal SC, where optimal gaze amplitudes ranged from about 30 to 100 deg.

#### 2.1 Network architecture

We modelled the SC gaze motor-map by a one-dimensional two-layer spiking neural network with a cortical input layer, and a layer of SC output neurons with the Brian2 spiking neural network simulator^[15]^. Each layer consists of 200 neurons, uniformly distributed on 0-5 mm, which corresponds to the SC gaze-motor map midline (horizontal saccade direction). The neurons in the input layer all had identical biophysical properties and transformed an externally applied input current into a Gaussian population of spiking activity, which is passed on to the SC neurons through one-to-one, topography-preserving, synaptic connections. For simplicity, the neurons in the input layer were assumed not to interact with each other. Furthermore, the spatial-temporal properties of the input current profile were assumed to be invariant for different gaze amplitudes (i.e., the same for all sites in the input layer).

The SC neurons process the input neurons’ spikes through their topographically varying intrinsic properties, as described in our earlier work^[16,17]^. In short, the biophysical parameters of the SC neurons, such as their adaptation time constant, their synaptic connection strengths with the input layer, and their lateral excitatory-inhibitory connections, were assumed to depend in a specific way on their location in the motor map and, as a result, identical spiking activities that arise from the input layer at different locations for different saccade amplitudes will lead to dissimilar responses of the SC cells (e.g., Fig. 4A-C).

The neural network’s output represents the desired gaze-shift amplitude by adapting the linear dynamic ensemble-coding scheme for the recruited population as proposed for eye-only saccades^[6,7]^ to gaze shifts. Thus, each single spike from neuron, *n*, in the population is assumed to contribute a small incremental movement, *m_n_*, called the neuron’s ‘spike gaze vector’. In our extended linear ensemble-coding model, the desired gaze trajectory is thus determined by a dynamic cumulative summation of all spike gaze vectors in the neural population:

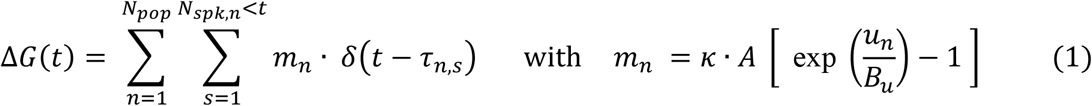

in which *t* is the current time, *δ*(*t* – *τ_n,s_*) represents a spike of neuron *n* at time *τ_n,s_, N_pop_* is the total number of active neurons in the population, and *N_spk,n_* <*t* is the total number of spikes fired by neuron *n* up to time *t*. The spike gaze vector depends exclusively on its rostral-caudal coordinate, *u_n_*, with *u_n_* ∈ [0 – 5] mm encoding the horizontal gaze-saccade amplitude in deg, and *κ* is a fixed scaling factor that only depends on the assumed cell density (in number neurons/mm). This scaling factor was calibrated for a horizontal saccade of 15 deg. The SC afferent mapping parameters (here: *A*=3.0 deg, and *B_u_*=1.4 mm) were adopted from monkey microstimulation data^[9,10,18]^.

Neurons in the model were described by the adaptive exponential integrate-and-fire (AdEx) model^[19,20]^ which is a conductance-based model with an exponential membrane potential dependence that can yield a variety of bursting dynamics with pnly few free parameters. Details of this neural model, including the chosen parameter values, are provided in the Supporting Information (Table S1), and in our previous work^[16,17]^. Here, we only highlight the major differences with the earlier published models.

#### 2.2 Saccade target representation by external input current

We provided a desired target vector, T, to the network by an external input current evoking population activity centered around the image point, *u_T_* (Eqn. 2). The central neuron in the input population receives the maximum input activation current, *I_0_*(*t*), while the other neurons in the input layer are stimulated by current strengths that decay as a Gaussian with distance from *u_T_*. The spatial-temporal external input current was thus described by a separable spatial-temporal function on the input neurons by:

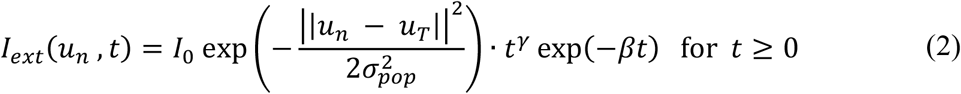

where *u_n_* is the anatomical position of a neuron in the input map, *σ_pop_* determines the size of the recruited input population, *t* is time (in s), *u_n_* is the location of neuron *n* (mm), and *I_0_* is the maximum input amplitude (pA). The time-dependent term is a gamma function, characterized by γ (skewness, dimensionless) and β (measure for the inverse input duration, in s^−1^).

#### 2.3 Superior Colliculus cells and influence of initial eye-in-head position

The neurons in the SC layer are characterized by several biophysical parameters, such as their adaptation time constant, their receiving lateral excitatory-inhibitory connections, and their synaptic input connections with the input layer, which all depend on their location in the motor map^[16,17]^. Each SC neuron thus receives a total synaptic input current, which is given by the excitatory synaptic weighted sum of the spikes coming from the sending neuron in the input-layer, and from all other SC neurons, whereby the latter is relayed by conductance-based lateral excitatory-inhibitory synapses (described by Eqn. S4, in Supporting Information).

We have shown previously^[16,17]^ that the adaptation time constant (τ_q_) systematically affects the peak-firing rate and burst-duration of the SC neurons, while the synaptic projection strengths between the input layer and the SC layer mainly affect the SC peak firing rate.

Neurons in the motor map communicate with each other through lateral interactions, organized according to a soft winner-take-all profile that is described by short-range lateral excitatory and long-range lateral inhibitory connections. As a result of this organization, the central cell in the neural population (the ‘winner’) synchronizes all other bursts in the population^[7,16]^. As the adaptation time constant and synaptic connection strengths between the two layers systematically decrease from the rostral to the caudal pole of the SC motor map, rostral neurons generate small saccades with high-frequency, short-lasting bursts of activity for their preferred saccade, while neurons at caudal sites, associated with large saccades, have lower peak firing rates and longer burst durations (see, e.g. Fig. 4). In line with neural recordings, however, the total number of spikes from the bursts in the population is invariant to the saccade amplitude, and even to the saccade kinematics: slow and fast saccades of the same amplitude are associated with different firing rates and burst durations but are encoded by the same number of spikes^[6,7]^. This invariance was achieved in the model by co-tuning the adaptation time constant (τ_q_) and lateral connection parameters^[16,17]^.

To extend the model to the control of eye-head gaze shifts, we included a modulatory signal proportional to the initial eye-in-head orientation that influences the firing properties of the SC neurons, with only a minimum effect on the total number of population spikes. As observed in single-unit recordings^[8]^, ipsilateral eye positions with respect to the target direction lead to a decrease in the neural peak-firing rate and an increase in burst duration compared to the straight-ahead eye direction, whereas a contralateral eye position leads to the opposite effect: higher peak firing rates with shorter burst durations (see Introduction).

Because the lateral connection strengths and adaptation time constant both affect the SC burst characteristics, we initially set out to modulate both parameters independently as function of eye position, by α_τ_(E_0_) and α_w_(E_0_), respectively. We found, however, that optimal tuning of the two parameters led to a strong mutual correlation (see Supporting Information, Fig. S1). We therefore lumped the two gains into a single modulation gain, denoted by α(E_0_). Best results were obtained for asimple near-linear relationship (Fig. 2), according to:

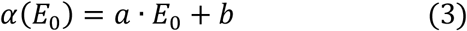

**Figure 2.**
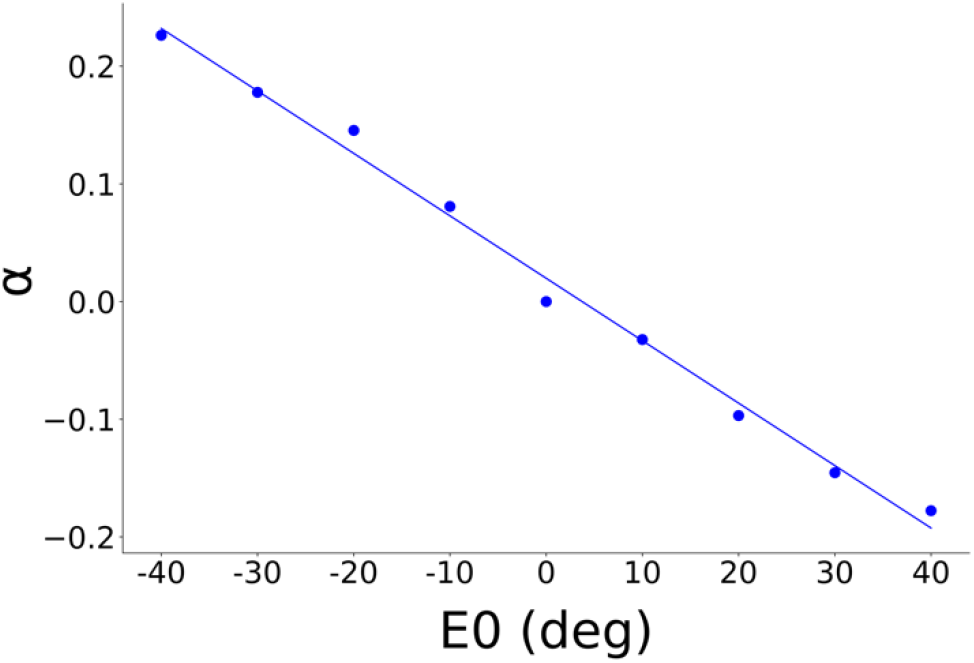
Optimal eye-position dependent gain, α, as used in the network simulations to modulate the neural activity patterns with the imposed constraints on firing rate and number of spikes. Best-fit result of Eqn. 3: a = −0.005 deg^−1^ and b =+0.018.

Note that the *α*(*E*_0_) was taken the *same* for all SC neurons. Thus, the eye-position signal was assumed to be distributed uniformly across the motor map. The network tuning for the adaptation time constant *τ_n_* as function of the map coordinate, *n*, and gain, α, is then summarized by:

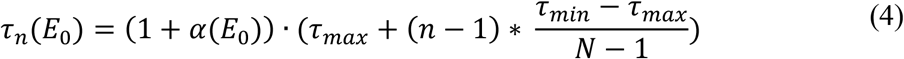

with N=200 the total number of neurons in the SC layer, *n* the neuron number, and [τ_min_, τ_max_] were set at [30, 60] ms, respectively.

Effectively, SC neurons received both excitatory and inhibitory potentials from cells endowed with different adaptation time constants, firing rates, and reversal potentials through the lateral connections (Supporting Information, Table S1). We modeled the latter by a Mexican hat-type connection scheme, where the net synaptic effect is given by the difference between two Gaussians^[16,17,20]^, but now each modulated by the same eye-position dependent gain:

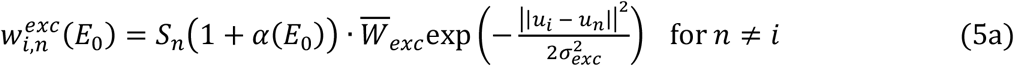

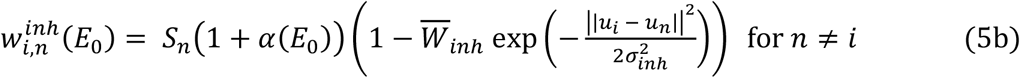

where 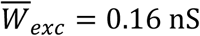 and 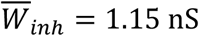 are fixed excitatory and inhibitory weight parameters. As explained in our previous work^[16,17]^, *S_n_* is the map-location-dependent gain, which makes the lateral interaction scheme site-dependent (see Eqn. S7 for details).

#### 2.4 Network tuning

We employed brute-force search algorithms to find suitable values for the gains of the adaptation time constant (Eqn. 4) and the lateral inhibitory and excitatory weight parameters (Eqns. 5a,b), the intrinsic properties of the AdEx equations of the SC neurons (Eqns. S1a, S2b) and the feedforward projection strengths from input to output layer 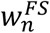 (Eqn. S3). The intrinsic biophysical parameters of the AdEx model (Supporting Information, Table S1) were optimized by systematically varying τ_q,n_, in combination with 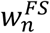 in a linear way with anatomical rostral-caudal location. Optimal values of the biophysical parameters are identical to those in our previous study for the initial eye position at 0° ^[16,17]^.

#### 2.5 Desired eye-movement trajectories

The population activity in the SC motor map encoded desired eye-head gaze shifts by the dynamic linear ensemble-coding scheme for the one-dimensional efferent motor map (Eqn. 1). The resulting instantaneous desired gaze-displacement trajectory, *ΔG(t*), was interpolated with a Savitzky–Golay filter to compute smooth instantaneous desired gaze velocity profiles.

#### 2.6 Neural movement fields

To analyze the movement-field properties of the neurons in the SC layer, we counted for each neuron the cumulative number of spikes in the burst during all horizontal gaze shifts with amplitudes between 5 and 55 deg in 2 deg steps, and for three initial eye-in-head orientations (−20, 0, +20 deg). We first fitted the neurons’ *static* movement-field functions for gaze saccades, by adapting the original quantitative model of Ottes and colleagues^[9,10]^ for eye-only saccades to eye-head gaze shifts, and included a potential effect of the initial eye-in-head orientation on the number of spikes in the burst^[21]^. Thus, the total number of spikes, N, of an SC cell was described by:

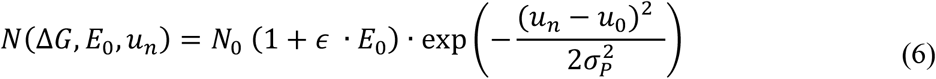

The static movement-field function predicts the total number of spikes in the SC burst to an arbitrary gaze shift for every cell *n* in the map and for each initial eye position. In our 1D model, the static movement field has four free parameters: N_0_ is the number of spikes in the burst for the neuron’s optimal saccade from straight ahead, u_0_ (in mm) is the optimal saccade coordinate in the SC motor map for the population, *ϵ* (in #spikes/deg) is the measured eye-position sensitivity of the neuron, and σ_p_ (in mm) quantifies the cell’s tuning width (spatial extent of the gaze-shift field as point-mapped on the motor map). Finally, *u_n_* is the anatomical SC coordinate of neuron *n* corresponding to the point image of its own optimal gaze shift, and is obtained by the (1D) afferent mapping function^[10]^:

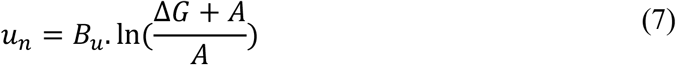

where B_u_ = 1.4 mm and A = 3.0 deg (see, e.g. Fig. 1B,E)^[9,10]^.

Next, we described the neuron’s firing profile by the *dynamic* movement-field function that predicts how the cumulative number of spikes in the burst of neuron *n* evolves as a function of time during each desired straight gaze-displacement^[6,8]^. We here extended our original concept for eye-only saccades^[6]^ by including the gain-influence of initial eye-in-head position, E_0_. As a consequence of the dynamic ensemble-coding model (Eqn. 1), the cell’s dynamic movement field is predicted to behave according to the following linear relation:

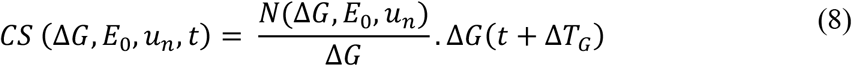

with Δ*G*(*t* + Δ*T_G_*) the desired straight trajectory (correcte for the fixed neural lead time of 10 ms), which increases monotonically from 0 to the final amplitude, Δ*G*. The time-independent factor in Eqn. 8, 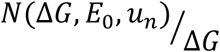, is the expected slope of the linear dynamic phase-relation between the ongoing (delayed) gaze shift and the expected cumulative number of spikes of the cell (see, e.g. Fig. 1D,F). This slope is proportional to the number of spikes from the static movement field (Eqn. 6), and is inversely related to the gaze-shift amplitude^[8]^.

#### 2.7 Sensorimotor transformation for generating eye-head gaze shifts

Based on our recorded behavioral and neurophysiological data^[8]^, we implemented a 1D sensorimotor model for the generation of eye-head gaze shifts, in which the SC cells act as a common gaze command to drive the eyes and head to the target location^[2,11]^. The population of recruited SC cells encodes the dynamic desired straight gaze trajectory, Δ*G(t*), through Eqn. (1)^[6,7]^. The parameters of the network cells are influenced by the distributed initial eye-position signal (Eqns. 3–5). Accordingly, the summed instantaneous firing rate of the population specifies an instantaneous desired gaze-velocity profile. The schematic outline of our computational model is shown in Fig. 3.

**Figure 3.**
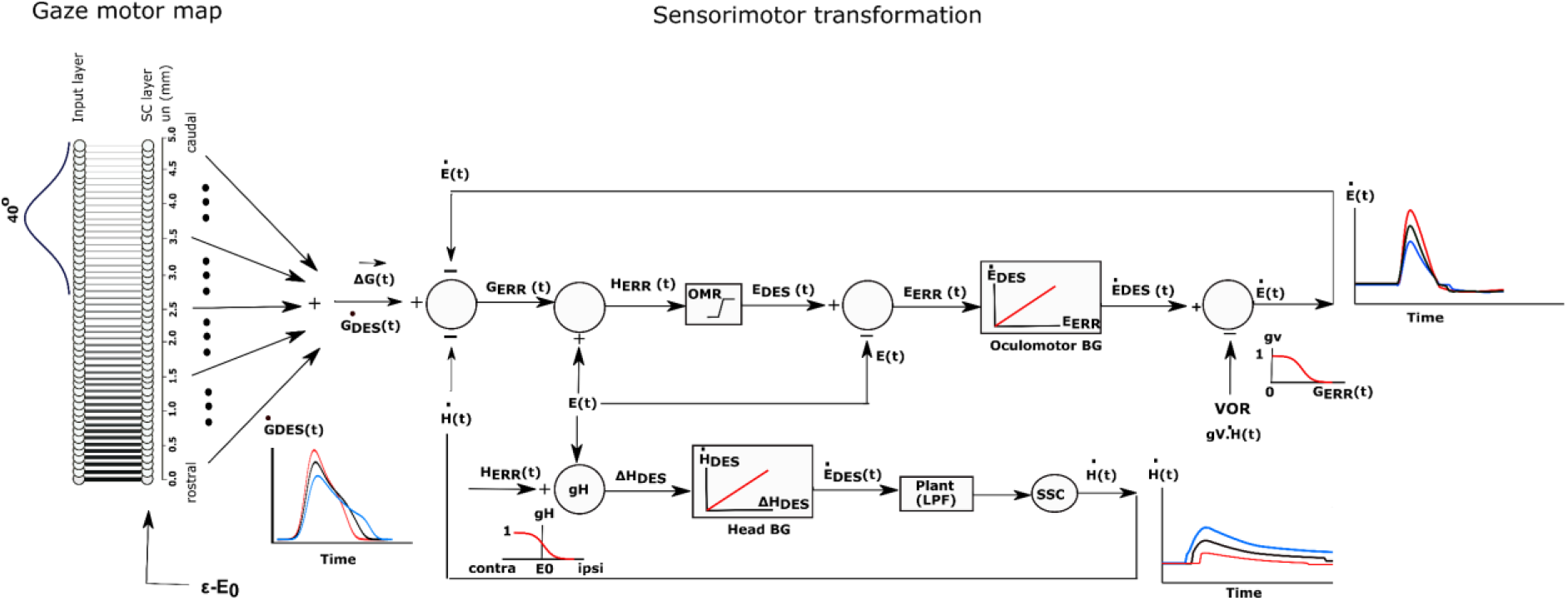
One-dimensional model for the generation of horizontal eye-head gaze shifts. The desired gaze-shift trajectory, ΔG(t), and its associated gaze-velocity profile, 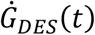, are encoded by the cumulative number of spikes (Eqn. 1) and the total instantaneous firing rate of the recruited SC population, respectively. Initial eye-in-head position, E_0_, modulates the SC cells’ firing profiles, as well as the contributions of the eye-and head movements to the gaze shift, ΔE, andΔH (inset lower-left) by influencing their relative timings. The vestibulo-ocular reflex (VOR) gain is modulated by the ongoing gaze error, G_ERR_(t) (inset, top-right). The upper part of the control scheme corresponds to the oculomotor system; the lower part to the head-motor system. The signal about initial and instantaneous eye position, E(t), is derived from the oculomotor neural integrator (not shown in the scheme). The comparator (left) subtracts and integrates the neural estimates of instantaneous eye- and head velocity from the total instaneous firing rate from the SC motor map, yielding the comman gaze-error motor command for the eye and head. See text, for further details.

#### 2.8 Model implementation

The firing rates of the population of recruited SC cells effectively encode a desired dynamic gaze-velocity signal, 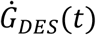 (by the summed instantaneous firing rates of all neurons). Inspired by Scudder’s model for eye-only saccades^[22]^, the instantaneous gaze-motor error then follows from the dynamic difference between the cumulative integral (spike count) of the desired gaze velocity and the true gaze velocity constructed from the oculomotor and head-motor burst controllers:

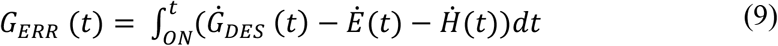

### Oculomotor system

Note, however, that the desired dynamic gaze-error signal may extend beyond the physical, head-centered, oculomotor range (upper section, Fig. 3). To prevent this from happening, the oculocentric gaze error of Eqn. 9 is therefore first transformed into a craniocentric (eye-in-head) error signal by adding the instantaneous neural estimate of eye position, which is obtained from the downstream oculomotor neural integrator^[23,24]^:

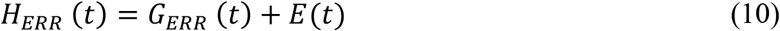

This dynamic head-centered error signal then drives both the oculomotor and head-motor control systems^[2,11]^. To keep eye position within the mechanical head-centered oculomotor range (OMR; here set between −30° and +30°), H_ERR_(t) for the eye was constrained by a soft limiter embodied by a hyperbolic tangent, yielding the actual desired horizontal eye-in-head position of the gaze shift to the target:

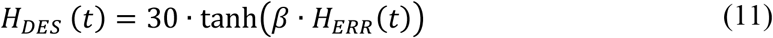

with β=0.03 deg^−1^. From this signal, the dynamic oculocentric eye motor-error is constructed:

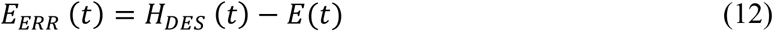

which drives the *linear* oculomotor burst generator, with a fixed gain of B_E_ = 60 s^−1^, to generate a desired eye-in-head velocity signal^[6–7]^:

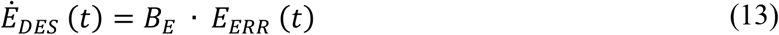

The neural estimate of the true eye-in-head velocity command for the oculomotor neurons, neural integrator, and eye plant (this final common oculomotor path^[23]^; not shown in Fig. 2, for simplicity) is obtained by linearly combining the desired eye-velocity signal with the gain-modulated vestibular-ocular reflex (VOR)^[25–27]^. The latter is driven by the true head velocity, and its gain, *g_V_*, is determined by the instantaneous gaze error:

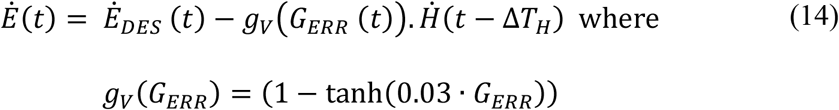

The VOR gain is close to 1.0 (i.e., is fully engaged) when the gaze error is small (the eye is on target), and it rapidly falls to zero for large gaze errors (e.g., at the start of the gaze shift)^[1,2,25]^.

### Head-motor system

In parallel, the head-motor system (lower section of Fig. 3) is directly driven by the head-motor error (Fig. 2), scaled by a gain that is assumed to depend on the initial eye position:

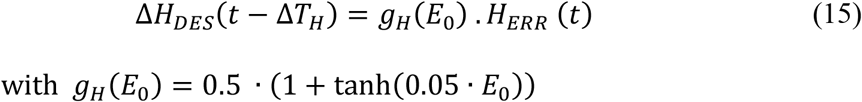

Importantly, the head-onset delay is assumed to depend directly on the initial eye orientation and on the desired gaze-shift amplitude. We here simply modeled this effect in a bi-linear way:

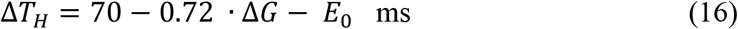

Thus, the delay increased for contralateral eye positions (E_0_<0) and decreased for larger gaze shifts and ipsilateral eye positions. The desired head velocity was subsequently generated by a linear head-burst generator with a fixed gain of B_H_ = 20 s^−1^:

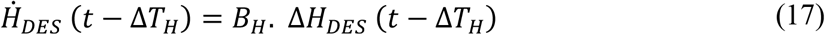

The actual head velocity, as measured by the vestibular canals, follows after passing the desired head-velocity command through a sluggish first-order low-pass filter as a simplified model for the head-motor plant:

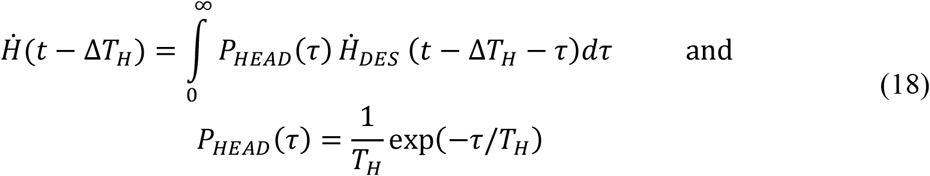

where we took T_H_ = 250 ms.

In the simulations, presented below, we varied the initial eye orientation *E_0_* between [-40°, +40°] and gaze-shift amplitudes Δ*G* between 1° and 60°.

## 3. Results

### 3.1 Eye-position influence on SC activity

Figure 4 shows the burst profiles for three example SC cells recruited for their optimal gaze shifts with the eye in nine different initial positions, varying from the contralateral (reddish lines) to the ipsilateral (bluish lines) side of the head from −40 to +40 deg in 10 deg steps. Note the systematic increase of burst duration and the corresponding decrease of the peak firing rate (by about 20%) as the eye-in-head position varies from the contralateral to the ipsilateral side.

**Figure 4.**
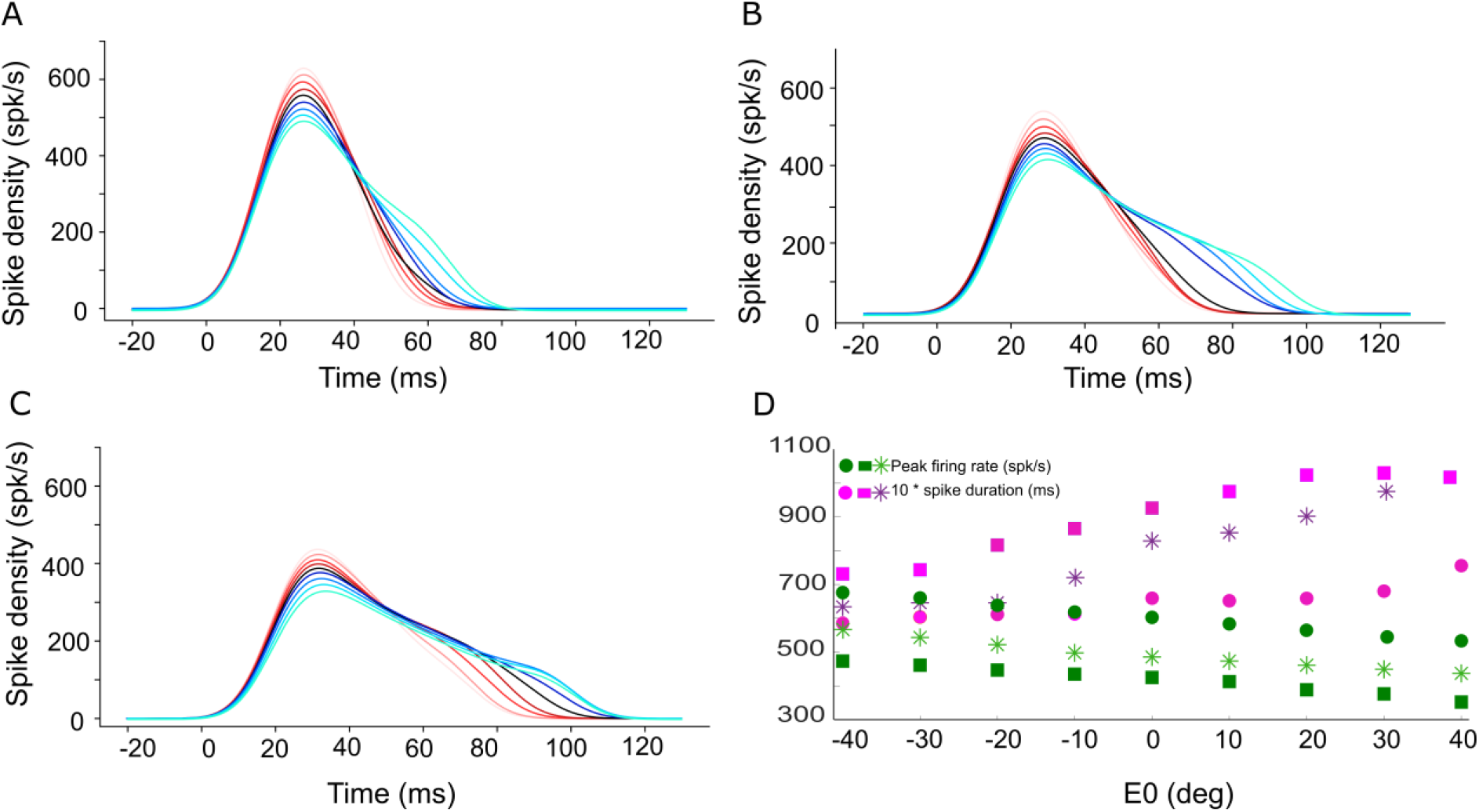
Instantaneous firing rate of the central neuron in the neural population for (A) 15 °, (B) 30 °, and (C) 45 ° gaze shifts with the initial eye-in-head position ranging from [-40,+40]^()^ (red to blue colors, respectively). The SC bursts start approximately 20 ms after the start of input current (at t=0). (D) The peak firing rate of the central neuron (green circles for 15 °, green stars for 30 ° and green squares for 45 ° gaze shifts) decreases, while the burst duration (pink symbols) increases, when moving from the more rostral (A) to the more caudal (C) side, and when the eye moves from the contralateral to the ipsilateral side of the head.

To tune the spiking neural network, the adaptation time constant and excitatory-inhibitory lateral connections in SC layer were modulated by the eye-position gain α in such a way that the number of spikes in the burst, as well as the total number of spikes of the neural population remained approximately invariant for all gaze-shifts and all initial eye positions. The result of this tuning is shown in Fig. 2. Figure 5A shows that the central cells in the three different neural populations indeed emitted approximately the same number of spikes (*N_spk,n_* ≈ 20) for the three desired gaze shifts (different symbols), and for all eye-in-head orientations, despite the substantial changes in the peak firing rates and burst durations of their firing profiles (Fig. 4). Similarly, the total number of spikes of the three neural populations remained practically invariant at about 450 spikes for the different gaze-shift amplitudes across all eye-in-head positions (Fig. 5B).

**Figure 5.**
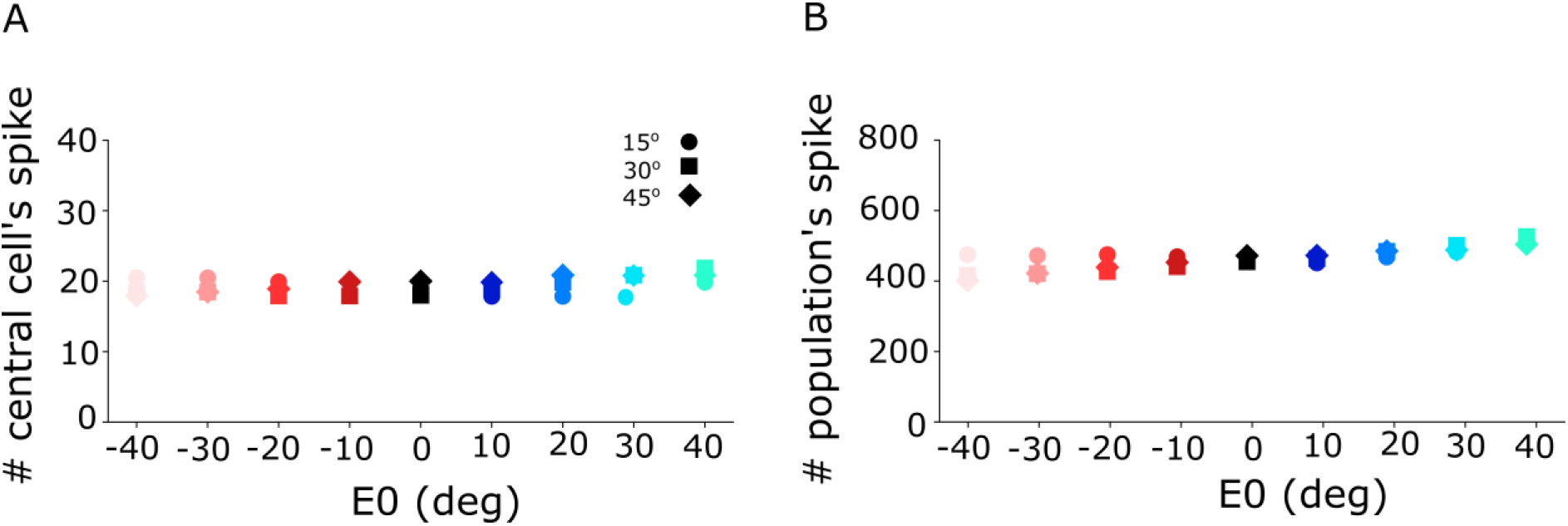
A) Number of spikes emitted by the central cell in the neural population varies only little between 18 and 21 spikes for different initial eye positions and for the three different gaze-shift amplitudes at 15 deg, 30 deg and 45 deg. B) The total number of spikes sent by the SC neural population to the downstream gaze-control circuitry remained virtually invariant across the different initial eye positions and gaze-shift amplitudes.

Figure 6 shows how the planned peak gaze velocity (given by the total summed peak firing rate of the population) increases with the desired gaze-shift amplitude (from Eqn. 1), and the effects on this desired velocity from the changes in initial eye-in-head orientation. The planned kinematics show the typical saturating relationship observed for head-restrained ocular saccades^[5–7]^, with a relatively slight modulation of the peak velocity because of the eye-position gain (Eqn. 3). Note, however, that in the case of eye-head gaze shifts, these planned gaze kinematics may become dissociated from the true gaze kinematics, because the SC neurons in the model do not sense any of the associated changes in the eye- vs. head-movement contributions that are determined in the downstream circuitry of the model.

**Figure 6.**
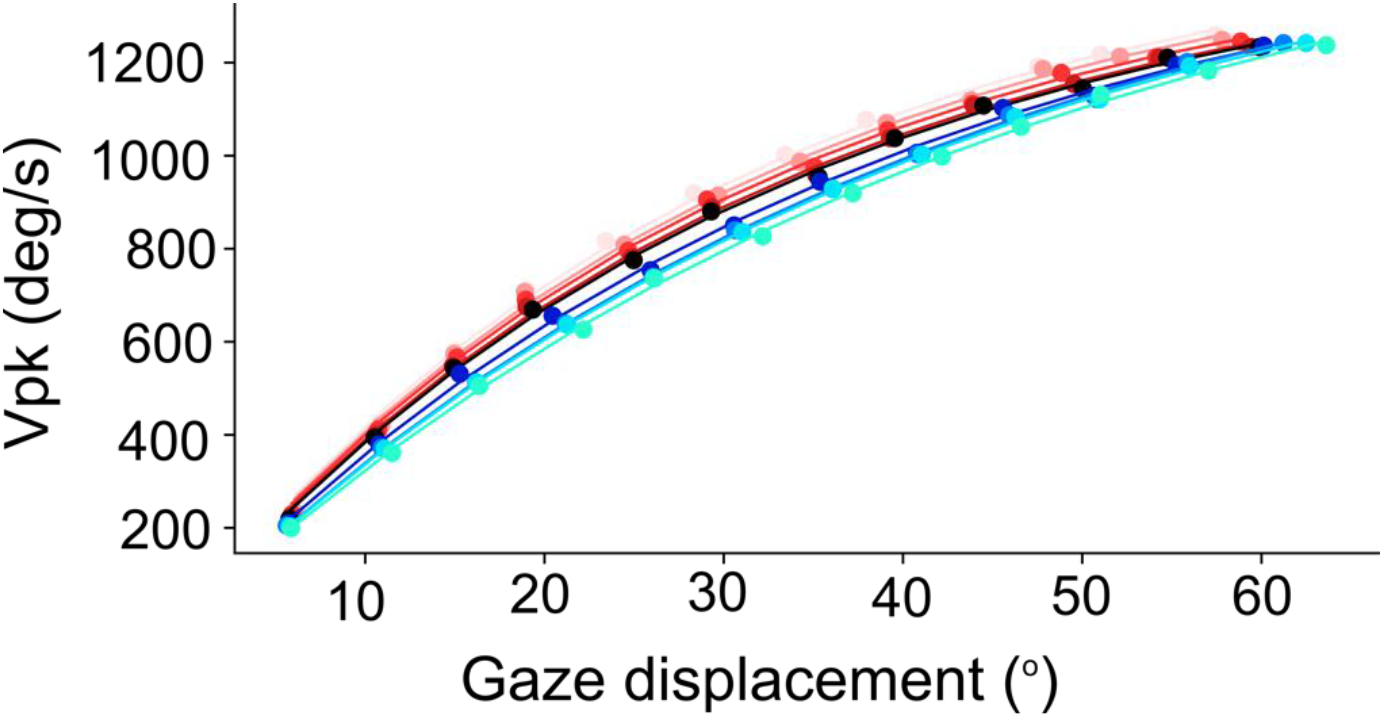
Nonlinear, saturating, increase of the planned peak gaze-velocity as function of gaze-shift amplitude as encoded by the SC population. Results are shown for nine initial eye positions (−40:10:40; different colors) and for 12 sites along the horizontal meridian of the motor map (dots).

### 3.2 Eye-head coordination

A horizontal saccadic eye-head gaze shift (ΔG) is composed of the linear sum of the instantaneous eye-in-head and head-on-neck orientations: Δ*G(t) = ΔE(t) + ΔH(t*). The contribution of the head movement to the gaze shift has a strong effect on the gaze kinematics. This point is illustrated in Fig. 6, which exemplifies two gaze shifts: one of 35 deg (Fig. 7A,C), and one of 55 deg (Fig. 7B,D).

**Figure 7.**
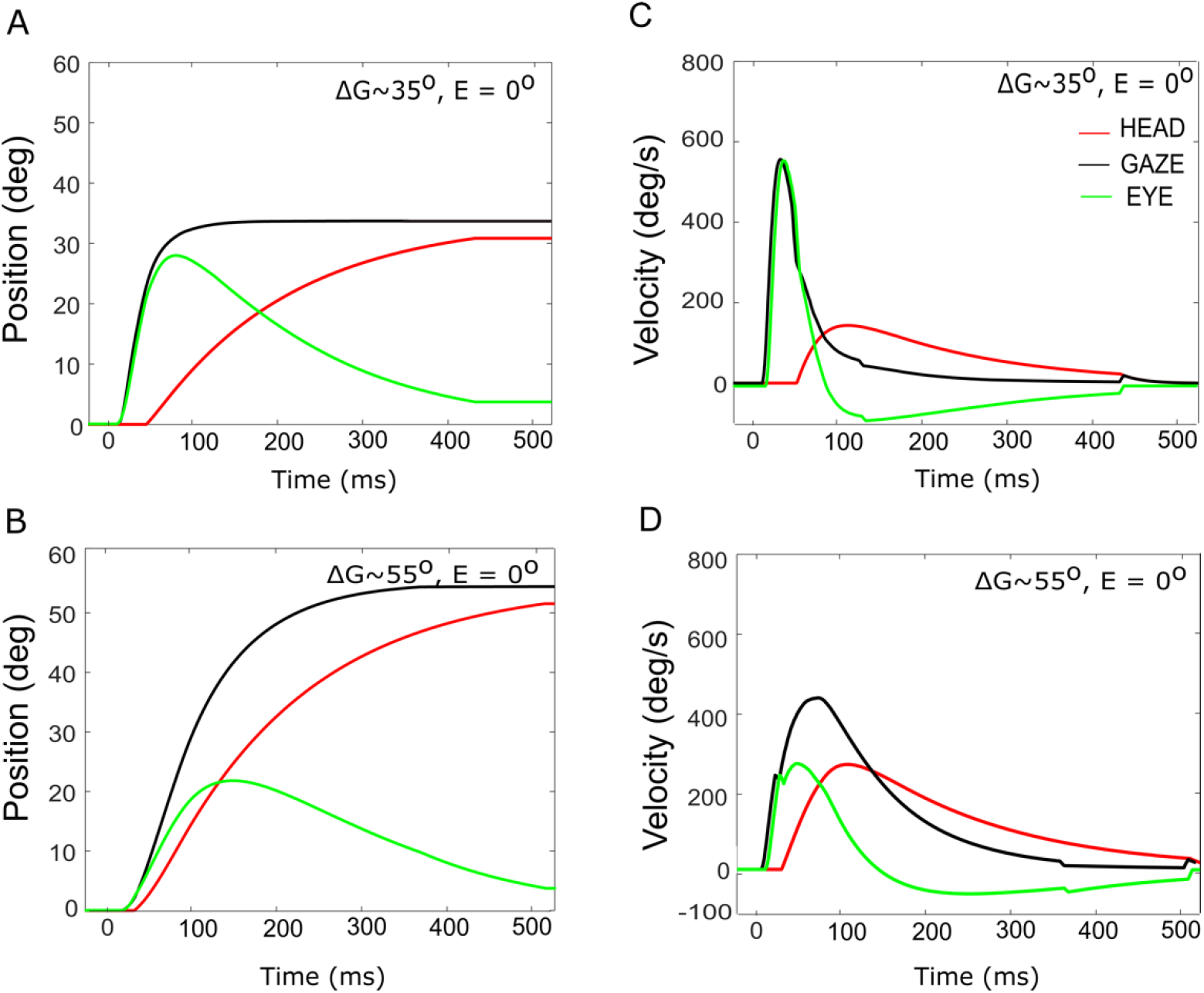
Gaze- (black), head- (red) and eye- (green) displacement (left) and associated velocity profiles (right) for gaze shifts with an amplitude of 35 ° (A,C) vs. 55° (B,D). Note that for the 55° gaze shift, the head onset is earlier, and the head contribution is considerably larger (about 20 ° vs. 5 °), causing the overall gaze velocity to drop considerably, such that it is even slower than the smaller gaze shift. The time axis is referred to burst onset in the SC (20 ms after the input). The additional delay of 10 ms of the gaze shift accounts for the efferent delays in the motor system.

Both gaze shifts start with the eye and head directed at straight ahead (i.e., E_0_=H_0_=0 °). For the larger gaze saccade, the head movement starts earlier after the eye-movement onset than for the smaller gaze saccade (Eqn. 16). Quite remarkably, because of the much larger head contribution in the latter case (about 20° vs. 5° at gaze offset), the 55° gaze shift is considerably slower than the 35° gaze shift, and therefore breaks with the well-known stereotyped main-sequence relationship for eye saccades^[5]^.

The influence of initial eye position on the kinematics of simulated gaze shifts, and on the associated head- and eye movements is further illustrated in Fig. 8, for fixed-amplitude 40° gaze saccades, generated from three initial eye-in-head orientations: contralateral (Fig. 8A), aligned (Fig. 8B), and ipsilateral (Fig. 8C). Note the different head-movement contributions during the gaze shifts and the associated changes in the gaze-velocity profiles for the different initial conditions. As initial eye-in-head position moves from contralateral (A) to ipsilateral (C), the relative eye movement gets smaller and slower and the head contribution to the gaze shift increases because it starts earlier. As a consequence, the gaze shift becomes slower^[1–3]^.

**Figure 8.**
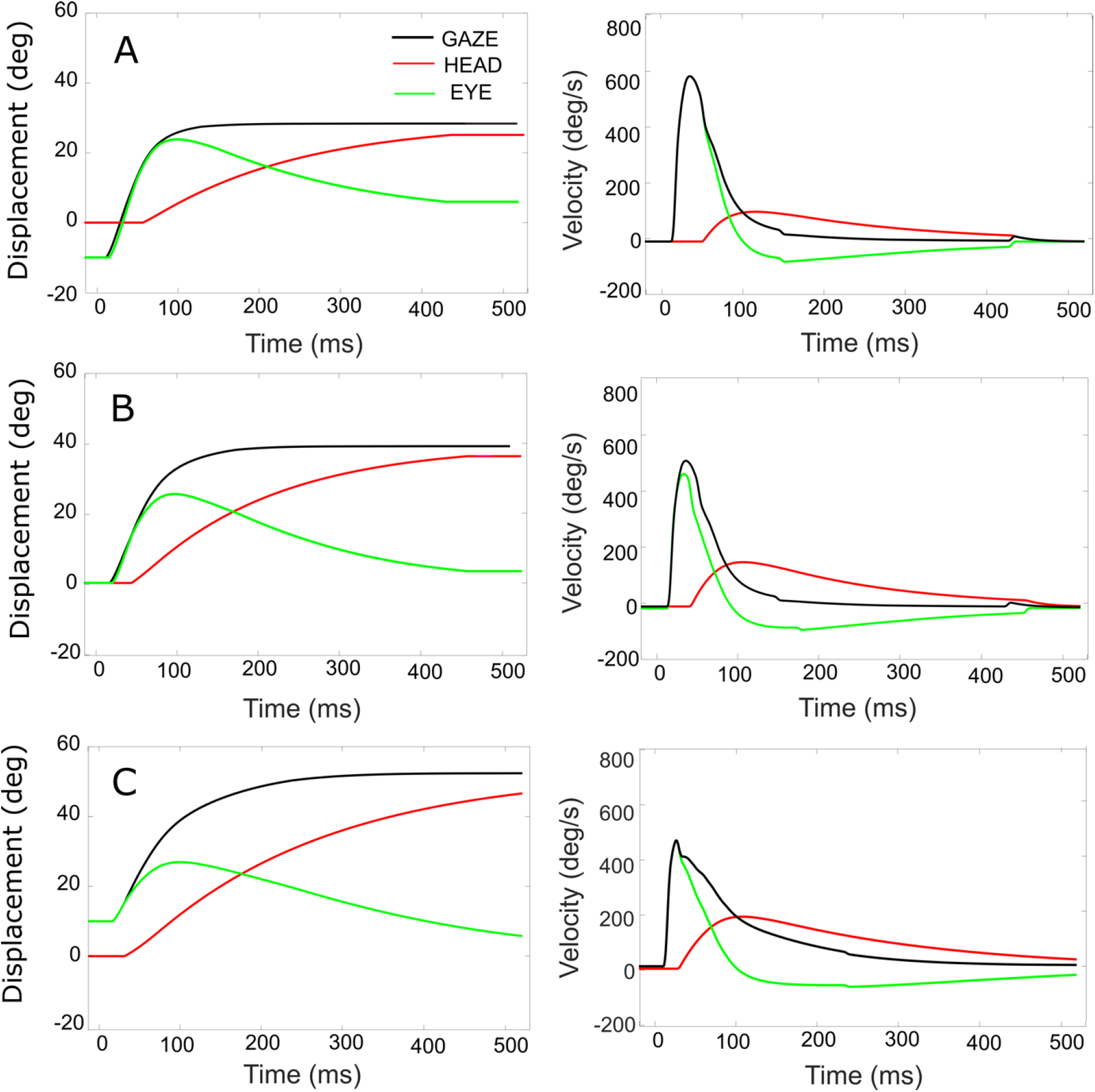
Simulated rightward gaze shifts of 40° amplitude, for three different initial eye-in-head fixations. A: Contralateral initial fixation, at E_0_=-10°. B: eyes at E_0_=0. C: Ipsilateral initial fixation, at E_0_ =+10°. Left-hand column: eye, head, and gaze trajectories. Right-hand column: eye, head, and gaze-velocities. Note the strong eye-position dependence of the contributions of eye and head to the gaze shift, as well as to the gaze kinematics and head-onset delay. The fastest gaze shift, with the largest eye movement, and smallest and latest head movement is obtained for the eye in the contralateral initial position (A). The slowest gaze shift with the smallest eye movement and the largest head movement is obtained for the eye in the ipsilateral direction (C).

The main-sequence relation for the model’s gaze shift (amplitude vs. peak-velocity) for gaze amplitudes between 5° and 60° is shown in Fig. 9A for five different eye-in-head conditions: aligned (E_0_ = 0; black), eye contralateral (at −10°, or −30°; red and brown symbols), and eye ipsilateral (+10°, +30°; dark and light blue symbols) of the target. Note that for small gaze shifts (Δ*G*<40 °), gaze velocity systematically increases with amplitude, just like for head-fixed eye saccades. However, for larger gaze shifts (Δ*G* >40 °), the peak gaze velocity starts to drop considerably with increasing gaze-shift amplitude. This effect was highly significant for both the contralateral and centered eye positions, and less strong for the ipsilateral eye orientations. Interestingly, a similar behavior was reported for monkey gaze shifts^[8,27]^. Figure 9B shows a representative example of a recording taken from a head-unrestrained monkey making large horizontal gaze shifts from three different initial eye orientations.

**Figure 9.**
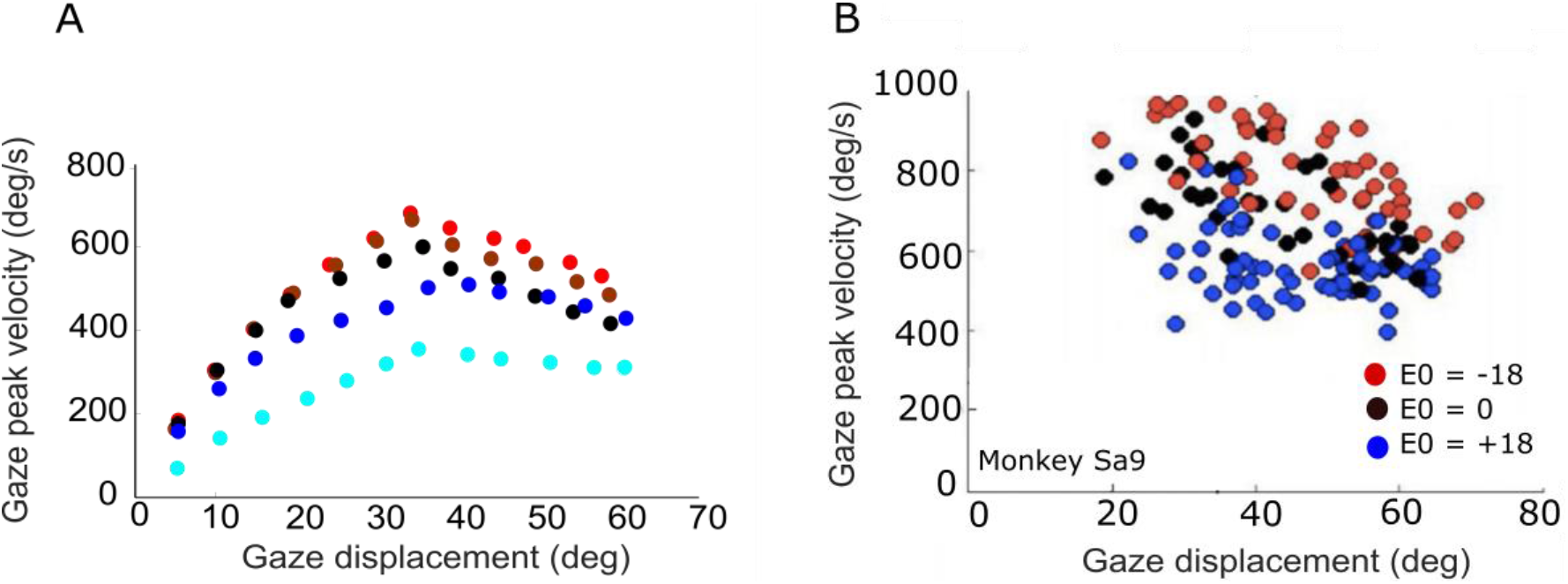
A) Peak gaze velocity as a function of gaze-shift amplitude for five different initial eye-in-head fixations (red (contra): −30 °, −10 °; black (center): 0 °; blue (ipsi): + 10 °,+30 °). Fastest gaze shifts occur when the eyes fixate contralateral to the gaze shift. Gaze shifts become markedly slower when the eyes fixate ipsilaterally (blue). B) Experimental data from large horizontal gaze shifts between about 20 and 70°, recorded from a macaque monkey for three different initial eye positions (−18°,0°,18°; same recording as in Fig. 1).

According to our model, this remarkable kinematic property is due to two factors: first, for large gaze shifts, the planned eye movement will readily approach the oculomotor range so that the eye-in-head velocity will start to plateau^[1–3]^. Second, at increasing gaze amplitudes the contribution of the (slower) head movement increases too (Fig. 7; Eqn. 15). As a result, the head velocity will increasingly dominate the gaze velocity profile when the gaze-shift amplitude increases. The effect is least pronounced for the far-ipsilateral condition because in that case the head trajectory already coincides nearly fully with the eye trajectory. As a result, the gaze shift is already close to maximally influenced by the head.

### 3.3 Static and dynamic SC movement fields

In this section, we analyze the static and dynamic movement-field properties of the model SC neurons for the different gaze-shift conditions shown above. This analysis links the the firing patterns of the spiking neural network (shown in Figs. 4–6) to the model’s output (eye- and head movement trajectories; Figs. 7–9).

As mentioned above (Fig. 6), the feedforward encoding of the gaze kinematics (by the population firing rate; Fig. 3) is expected to become dissociated from the actual gaze kinematics (Figs. 7–9) when the head starts to contribute to the gaze shift, since the SC neurons in our model have no access to a head-movement signal, which in turn strongly determines the gaze kinematics. Without a head movement, however, the dynamic linear ensemble-coding model of Eqn. 1 predicts a one-to-one relationship between the eye-saccade velocity and the population firing rate^[6]^, as well as the straight-line relations between the cumulative number of spikes and the instantaneous eye displacements (dynamic movement fields). Figure 10 illustrates these properties for 10° gaze shifts without a head movement (i.e., we set gH = 0) generated from three different initial eye positions.

**Figure 10.**
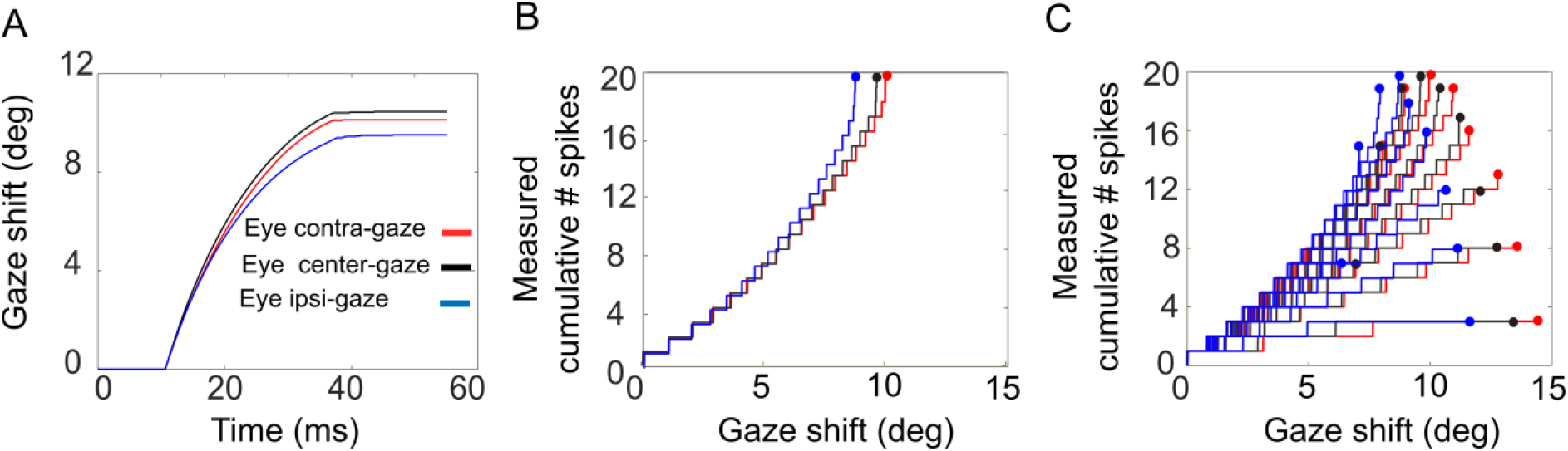
Gaze shifts of 10° amplitude without a head movement, for three different initial eye-in-head fixations: E_0_=-10° (red), 0° (black), +10° (blue). A) Gaze trajectories. Note the different speeds with varying eye position. B) Phase trajectories of the cumulative number of spikes as function of ongoing (back-shifted by 10 ms) gaze displacement. Note that the curves superimpose although the gaze kinematics differ substantially. C) Cumulative number of spikes in the burst vs. ongoing (back-shifted) gaze displacement for all saccades (7-15 deg) into the cell’s movement field (cf. Fig. 1D).

Although the gaze kinematics are considerably affected by the initial eye orientation (Fig. 10A), the phase curves that relate the cumulative number of spikes of the neuron to the instantaneous change in gaze for the optimal gaze amplitude (aligned by ΔT_G_ = 10 ms) are quite similar (Fig. 10B). This behavior is a direct consequence of the linear ensemble-coding scheme (Eqn. 1), in which each spike of each cell is assumed to contribute a fixed gaze displacement. Note, however, that since the ordinate represents the output of a single neuron, whereas the abscissa is the output of the total neural population, the tight resemblance of the three phase curves should be understood from the high level of synchronization of the bursts among all recruited cells. The latter is caused by the soft winner-take-all lateral interaction mechanism in the motor map^[16]^. Figure 10C shows the phase curves for all gaze saccades into the cell’s movement field for the three different initial eye positions. The dots at the end of the curves correspond to the final gaze displacement and total number of spikes (cf. with Fig. 1D). These dots should follow the static movement field of Eqn. 6, where the total number of spikes depends on gaze amplitude (between about 6 and 15 deg) and initial eye position. Note that the phase curves have different slopes for each gaze amplitude, like in Fig. 1D. For eye-only saccades, neural recordings have shown that each of these lines may be predicted by the dynamic movement-field description of Eqn. 8^[6]^. It is not a-priori obvious, however, that the same holds for our eye-head gaze-control system with its inherent nonlinearities (Fig. 3).

To look at this point in a little more detail, we analyzed the static and dynamic movement-field relationships of our model SC neurons during eye-head gaze shifts over the full range from 1-60 deg. The results for three representative neurons are shown in Fig. 11.

**Figure 11.**
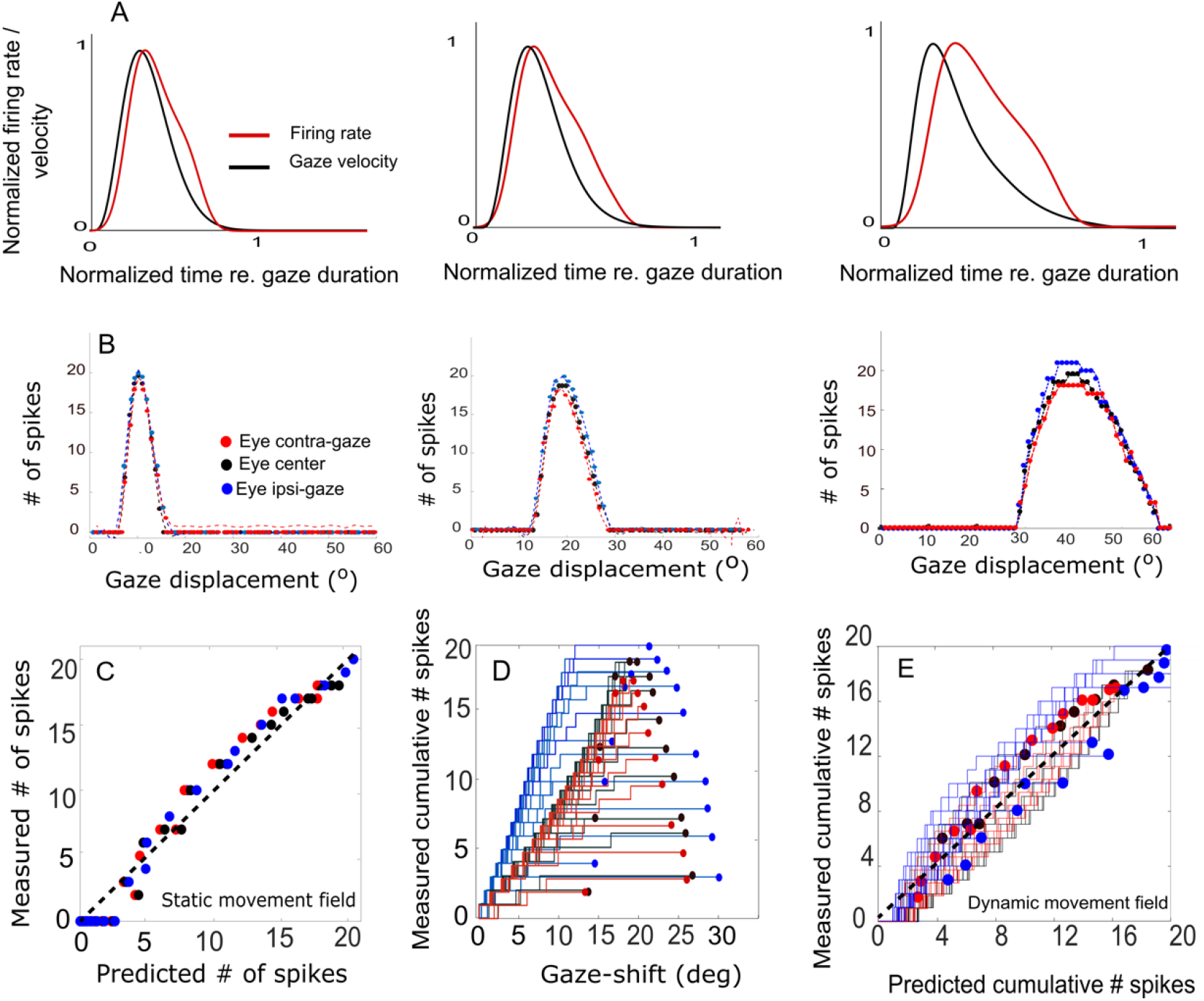
(A) Three example trials demonstrating the relation between the cell’s firing-rate profile and the instantaneous gaze velocity for three initial eye positions in a 20 deg gaze-shift. For ease of comparison, both variables were normalized to gaze duration and to their maxima. The correlation is highest for the contralateral situation (r=0.92) and decreases for the ipsilateral condition (r=0.69). (B) Plot of the three static movement fields for cells with their optimal saccades at 10 °, 20 °, and 40 °, respectively (Eqn. 6) for three different initial eye positions. Note that the movement fields vary little with the initial eye position. (C) The static gain-field model (Eq. 6) captures the data well for all gaze shifts and initial conditions for the cell with its optimum at 20 ° (see also Table 1). (D) Phase trajectories of the cumulative number of spikes of the cell as a function of ongoing gaze displacement along the straight gaze vector (dynamic movement field). (E) Test of the dynamic movement-field model of Eqn. 8 on the spike trains during all fast (red), intermediate (black), and slow (blue) gaze shifts into the movement field of the 20 deg cell. Compare with Fig. 1.

First, we examined how much a neuron’s firing rate correlates with the instantaneous gaze velocity (cf. Fig. 1C). Panel 11A shows the gaze-velocity traces (black) and instantaneous firing rates for the neuron encoding Δ*G*=20 deg (red) for a fixed 20° gaze shift from three initial eye positions. For illustrative reasons, the data traces were normalized with respect to their peak amplitudes and to the respective gaze duration (set at 1.0). The normalized traces appeared to correlate well, although for the ipsilateral condition the peaks of the two curves are no longer well aligned. Overall, we obtained correlations r>0.8 for all trials in the majority of the 200 model cells.

In panel 11B we analyzed the static movement-field properties of three cells for the three initial eye positions, by plotting the total number of spikes in the burst for each neuron as a function of gaze-shift amplitude. The neurons were located near the rostral, central and caudal sites of the SC gaze-motor map, respectively. By moving from rostral to caudal areas of the map, the movement field of the cells covers a larger range of gaze-shifts, which is caused by the compressive logarithmic afferent neural mapping^[9,10,18]^ (Eqn. 7). Thus, the 10 deg cell is recruited for gaze shifts between about 6 - 15 deg amplitude (range 9 deg), while the 40 deg neuron is involved in gaze shifts between 30 - 60 deg (range 30 deg). It can be seen that the three neurons emitted a fixed number of ~20 spikes for their optimal gaze shift and that the number of spikes varied only little (but systematically) across the different initial eye positions.

For example, for the 40 deg cell, with ε=0.0053 spks/deg and E_0_ = 20 deg, the expected maximum number spikes will increase from 20 to 22 spikes (as 20 · 0.0053 · 20 = 2.12). For ipsilateral eye orientations (E_0_>0) the number of spikes was indeed slightly higher than for contralateral eye positions. The dotted lines represent the fitted static movement-field curves through these data (Eqn. 6). The optimal fit parameters for the three cells are given in Table 1.

**Table 1:** Fitted movement-field parameters (Eqn. 6) for the three cells shown in Fig. 11B.

| Cell ($\Delta G$) | $u_n$ (mm) | $N_0$ (#spks) | $u_0$ (mm) | $\sigma_{pop}$ (mm) | $\varepsilon$ (#spks/deg) |
| --- | --- | --- | --- | --- | --- |
| 10 | 2.05 | 21 | 2.06 | 0.49 | 0.0017 |
| 20 | 2.85 | 19.83 | 2.85 | 0.47 | 0.0021 |
| 40 | 3.73 | 20.2 | 3.72 | 0.48 | 0.0053 |

Figure 11C shows the predicted number of spikes from the static movement-field model (Eqn. 6) for the 20 deg cell with its optimal parameters (Table 1; cf. Fig. 1D). The static movement-field model predicts the data quite well (*r* = 0.98).

Panel 11D presents the phase plots for the spike trains for the neuron with its optimal saccade at 20° (the central neuron in panel 11B). It shows the cumulative number of spikes, *CS(t*), as function of the dynamic gaze-shift vector, Δ*G(t+ΔT_G_*), with Δ*T_G_* = 10 ms. Note that each trajectory has a different slope, and endpoint that varied considerably for each gaze shift. Note also the influence of initial eye position on the cumulative number of spikes in the burst, which appears to organize the phase trajectories in three different clusters: blue for ipsilateral, black for central and red for contralateral initial eye positions.

To describe how the cumulative number of spikes evolves during the gaze displacement, we determined the prediction of the dynamic movement-field (Eqn. 8) to see how well it captures the variability in the cell’s spiking behavior. To that end, we used the static movement-field result from Eqn. 6 to the total spike counts to find the slope of the dynamic phase-relation. The result for all the data of panel 11D is shown in Fig. 11E. The overall correlation between the measured and predicted instantaneous spike counts is very high: r = 0.98.

## 4. Discussion

We constructed and tested a feedback control model for primate eye-head gaze shifts that was driven in a feedforward way by the output of a spiking neural network model of the midbrain SC. In our new computational model, the initial eye-in-head orientation influenced the dynamical characteristics of the SC cells in a uniform way, such that their bursting characteristics varied systematically with changes in eye position. We tuned the parameters of the cells such that the firing rates would monotonically increase with contralateral eye positions, and decrease for ipsilateral eye positions without appreciably affecting the total number of spikes in the bursts. In addition, initial eye position affected the contribution of the eye and head to the gaze shift by modulating the relative timings of the eye- and head-movement onsets. Depite its simplicity, the model produced horizontal eye-head gaze shifts with correct kinematic properties for a wide variety of initial conditions and amplitudes, together with neural response patterns in the SC motor map that faithfully resembled neurophysiological recordings from head-unrestrained monkeys.

### SC modulations

Our earlier work had indicated that the joint tuning of three biophysical parameters of the model SC neurons in the network determine both the firing rate and the number of spikes in the burst: the adaptation time constant, the top-down connections from the input layer, and the strength of the lateral intracollicular interactions. For simplicity, we here let the eye-position signal only affect the intrinsic SC parameters, i.e. *τ_q,n_* and the excitatory/inhibitory lateral connection strengths. Quite remarkably, in tuning the network for the imposed constraints we could obtain the required neural modulations with a single, simple linear gain control on both intrinsic variables (Eqns. 3–5, Fig. 2, and Fig. S1).

Alhough we did not attempt to optimally fit the neuronal firing patterns of our model neurons to those from real SC recordings in monkey (like in Fig. 1), the overall response behaviors of our model resulted to be quite similar. For example, although the aim was to modulate only the firing rate (and burst duration), but not the number of spikes, the cells nonetheless showed a small positive sensitivity on their number of spikes for eye position, since we found that the gainfield parameter ε > 0 for all neurons (e.g., Figs. 5 and 11; Table 1). This led to a small increase or decrease with one to two spikes for ipsilateral (slow movements) vs. contralateral (fast movements) eye positions and for small and large gaze shifts, respectively (note that ΔN = ε·N_0_E_0_). Interestingly, the eye-position sensitivities of our model neurons resulted to be very similar to values obtained from real neurons in monkey. For example, for the cell in Fig. 1 with Δ*G_opt_* = 37 deg we obtained ε=0.0024 spikes/deg, and in [8] we reported ε=0.0063 spks/deg for a cell with an optimal gaze vector of 57 deg amplitude.

Our model neurons yielded ε = 0.0017, 0.0021 and 0.0053 spikes/deg for the 10°, 20° and 40° gaze-shift neurons, respectively. This suggests that the modest eye-position sensitivity of the number of spikes increases with gaze-shift amplitude, even though the influence of the eye-position signal on the neurons was distributed homogeneously acoss the motor map. The underlying mechanism for this phenomenon can be explained by the position-dependent tuning of the relevant neuronal parameters across the map as follows: The number of spikes in the burst and the peak-firing rates are determined by the precise tuning of the adaptation time constant and lateral connection strengths^[16,17]^, which are all location dependent in our model. Because of this, the model produces high firing rates and short burst durations at rostral sites, vs. lower firing rates and longer burst durations at caudal sites, which underlies the nonlinear main-sequence of saccades^[5,6,7,16]^ (e.g., Fig. 6). As a result, however, the fixed gain effect of *E_0_* on the adaptation time constant (Eqn. 4) and lateral connections (Eqns. 5a,b) will differentially affect the bursting characteristics of these neurons too. For example, since *S_n_* ~ 1 – 0.04·*u_n_^2^* in Eqns. 5a,b the gain effect of *E_0_* will be stronger at rostral sites (*u_n_* small) than at caudal sites (*u_n_* large). Thus, the relative tuning of the relevant cell parameters becomes slightly imbalanced for non-zero eye positions, leading to a (small) effect on the number of spikes. As this effect will be site dependent, it will therefore vary with gaze amplitude.

Although it is tempting to speculate that a similar mechanism might apply to real SC neurons, there is no evidence to support it other than that it has been shown that the number of spikes in the SC burst for ocular saccades varies with eye-in-head position^[21]^. The neurobiological mechanism of this effect, however, remains elusive as extracellular recordings of spikes provide access to neither the intrinsic neuronal mechanisms, nor to the nature of the neuronal input. However, the simplified set of two coupled differential equations that determine the dynamics of our model neurons^[16,17]^ indicates that modulation of a single intrinsic parameter (the adaptation time constant) in combination with lateral feedback of spiking activity through modulated synaptic connections may suffice to produce the eye-position effect. Yet, it cannot be excluded that when using the full set of Hodgkin-Huxley equations to model the neurons, modulation of other neural parameters could lead to a similar performance.

### Gaze-shift kinematics

The stereotyped amplitude-peak velocity relationship for ocular saccades^[5]^ does not hold for eye-head gaze shifts (Fig. 9). First, variation of initial eye position strongly affects the gaze kinematics as it is a strong determinant for the contribution of the head during the gaze shift. In the model, this is achieved by a simple linear influence of *E_0_* on the onset delay of the head with respect to the eye. The earlier the head starts to move, the more time it has to influence the gaze-feedback loop by interacing with the eye-velocity signal. Thus, with the eye looking in the ipsilateral direction of the target (*ΔT_H_* reduced), the head contribution is substantially larger than when looking contralaterally, causing the latter to be faster than the former (Fig. 7). Also the planned gaze amplitude affected the head contribution by reducing the head-onset delay in proportion with ΔG (Eqn. 16). This latter effect, which functionally helps the system to overcome the limited oculomotor range to acquire the target, leads to the perhaps surprising fact that the peak gaze velocity starts to *decrease* for gaze amplitudes exceeding about 40 deg (Figs. 8, 9A). This phenomenon, which is strongest for the fastest gaze shifts (i.e., with the eye contralateral) is also clearly seen in monkey gaze-shift data (Fig. 9B)^[8]^. Note that this effect is not caused by the modulatory effects of eye position on the SC firing rates, which still encoded a monotonically increasing nonlinear desired gaze-velocity signal (Fig. 6).

According to the linear ensemble-coding model for eye-only saccades, the SC firing rates of the population directly encode the instantaneous eye velocity (and hence, the stereotyped nonlinear main-sequence relationship). This conclusion was based on the argument that simulating saccades with the recorded spike patterns from a large number of neurons through Eqn. 1 fully explained the trajectories and instantaneous kinematics of eye saccades, even though the entire brainstem model for the oculomotor system was made linear. The simple spike-count model also applies to slow saccades, e.g. when SC firing rates are reduced due to changes in initial eye position (Fig. 10), for saccades to remembered-targets^[28]^, or during blinks^[6]^, as in all these cases the total number of spikes in the burst remained virtually invariant.

The gaze-control model presented here, however, is no longer linear although the burst generators for the eye- and head motor systems were still modeled by simple linear input-output characteristics (Fig. 3). At least three nonlinearities play a role in the model that potentially break with the straightforward linear transfer characteristic of the oculomotor ensemble-coding model: (i) the eye-position and gaze-amplitude dependent delay of the head-movment onset, which affects the contribution of the slower head movement to gaze shifts; (ii) the nonlinear oculomotor range; (iii) the varying gain of the VOR. Yet, despite these nonlinearities, the relationships between the instantaneous cumulative number of spikes of individual neurons and the ongoing planned gaze displacement remained remarkably close to linear (Fig. 1D for a real recorded neuron, and Fig. 11D for our model neurons). As a result, the dynamic gaze-movement field function adequately described the neural tuning for instantaneous gaze shifts and their kinematics for a wide variety of initial conditions (Fig. 1F and Fig. 11E).

### Optimal control?

It has been suggested that the main-sequence relations for ocular saccades betray a speed-accuracy trade-off in combination with an undershoot strategy^[26]^ for the oculomotor system that optimally deals with the detrimental effects of multiplicative noise in neural control signals^[30–32]^ and uncertainty in the peripheral visual input. By reducing the high-frequency noisy impulse from the saccadic burst generator on the eye muscles for large saccades, the system would thus avoid the danger of saccadic overshoots that would further increase the total time for the fovea to acquire the target^[29]^. In our earlier work, we have argued that such a strategy would be best embedded at a level where signals are still encoded in an omnidirectional abstract vectorial format, rather than at the level of (much more complex, high-dimensional) individual muscle-control signals, and that the SC motor map could be an excellent candidate for such an optimal control principle^[7]^. The tight synchronization of the saccade-related bursts within the population, in combination with the apparent encoding of the saccade kinematics at the level of the motor map, seems to support this notion^[7]^. Indeed, simulating saccades with neural data applied to Eqn. 1 produced all the kinematic features and straight trajectories seen in real saccades, without having to resort to an ad-hoc saturating nonlinearity and component cross-coupling schemes in the brainstem burst generators^[6]^.

In line with this, it stands to reason that also eye-head gaze shifts would follow an optimal control strategy, albeit that the cost function to be minimized may differ from eye-only saccades. The oculomotor system only needs to worry about speed (time to target) and accuracy (foveation), whereby energy expenditure would be of minimal importance as the eye has negligible mass. This is not true for the head, and therefore a metabolic cost (e.g., total kinetic energy expenditure) might have to be included in the total movement cost as well^[33–35]^.

In light of this idea, what could be the role of the eye-position signal in the SC? We here speculate that the combined modulatory influence of eye position on the SC firing rates and on the head-onset delay might be an adaptation of the system to optimize the optimal control costs for combined eye-head gaze shifts. As the head’s moment of inertia is considerable, and hence it’s initial acceleration rather slow when compared to the eye, the system aims to minimize the contribution of the head to the gaze shift to reduce metabolic costs and at the same time optimize speed. However, because of the limited eye-in-head oculomotor range, significant head movements unavoidably need to be planned for all gaze shifts exceeding about 20 deg when the eyes don’t look in the contralateral direction. The uncertainty (i.e., intrinsic noise) in head-movement control signals is likely to be higher than for the eye, as the latter will hardly ever be influenced by external loads or forces, and has relatively simple plant mechanics (only rotations). As a consequence, and in line with speed-accuracy trade-off, the central command from the SC should account for the additional noise in its gaze-control signals when large head movements are needed. This would be achieved by lowering the SC firing rates (affecting speed and energy use) without (appreciably) changing the total number of spikes (which, in our model, determines gaze-shift accuracy).

### Limitations and future work

The one-dimensional spiking neural network model presented here can only generate horizontal gaze shifts. Although computationally more costly^[36]^, it is relatively straightforward to extend the SC motor map to a two-dimensional spiking neural network that enables the programming of eye-head gaze shifts in all directions.

It would then be interesting to also extend the downstream gaze-controllers to the full repertoire of 3D eye-head gaze shifts (i.e., horizontal, vertical, cyclotorsional). Eye-only saccades without head movements follow the well-known kinematic constraint of Listing’s law^[37]^, which states that all voluntary saccades are programmed as single-axis rotations whereby the axis of rotation ensures that the 3D orientation of the eye has zero cyclotorsion throughout the trajectory. We have recently developed a 3D model of the eye that closely followed Listing’s law and produced realistic main-sequence relationships and component cross-coupling for oblique saccades in all directions by applying optimal control of the physical model that minimized the total cost of speed, accuracy, and total force exerted by the six extraocular muscles on the eye during peripheral fixations^[38]^.

Listing’s law, however, does not hold for head-unrestrained gaze shifts^[39]^, not for the eye-in-head, the eye-in-space, or for the head itself. Instead, the involved motor systems are constrained by Donders’ law, which states that each 3D orientation (of eye and head and, consequently, also gaze) has a unique cyclo-torsional state, independent of the trajectory that brought it there. One reason for this is the involvement of the VOR towards the end of (and after) the gaze shift (e.g., Figs. 6 and 7), which requires the full three degrees of freedom needed to compensate the eye movement against any change in the ongoing 3D head orientation. Behavioral experiments have demonstrated that under certain initial conditions the eye-in-head can thus even obtain a cyclotorsional angle of about 15 deg during an eye-head gaze shift^[40]^. The underlying neural control strategies for such movements are highly nontrivial, and also require detailed knowledge of the 3D kinematics and dynamics of the eye-^[38]^ and head motor plants.

Although powerful computional models have been proposed also for 3D eye-head gaze shifts^[41,42]^, so far none of these models have incorporated the putative role of the SC in 3D gaze control. Presumably, the SC motor map issues a 2D (horizontal/vertical) desired gaze-displacement vector^[43,44]^ to the brainstem – cerebellar – skeletal motor systems, from which the appropriate 3D dynamic control signals will have to be derived. Our future work will aim to extend the current 1D model of Fig. 3 to a full 3D gaze-control system.

## Acknowledgments

This work was supported by European Union Horizon 2020 Programme, ERC Advanced Grant (2016) “Orient” 693400 (to AA and AJVO). The authors greatly acknowledge the kind and generous hospitality of dr E Freedman (School of Medicine, Univ of Rochester, NY, USA) to host AJVO on a sabattical stay in the lab in order to collect the neurophysiological and behavioral recordings from monkeys S and P, and to dr S Quessy, dr J Quinet, and dr M Walton for their kind and expert help in collecting the monkey data and many fruitful discussions.

## Author Contributions

AA and AJVO analyzed the data and interpreted the results of the experiments; AA prepared the figures and drafted the manuscript; AA and AJVO edited and revised the manuscript, and approved the final version of manuscript; AJVO conceived and designed the research.

## References

1. Guitton D, Volle M. Gaze control in humans: eye-head coordination during orienting movements to targets within and beyond the oculomotor range. J Neurophysiol. 58(3):427–459, 1987

2. Goossens HH, Van Opstal AJ. Human eye-head coordination in two dimensions under different sensorimotor conditions. Exp Brain Res. 114(3):542–560, 1997.

3. Freedman EG, Sparks DL. Coordination of the eyes and head: movement kinematics. Exp Brain Res. 131(1):22–32, 2000.

4. Kardamakis AA, Grantyn A, Moschovakis AK. Neural network simulations of the primate oculomotor system. V. Eye-head gaze shifts. Biol Cybern. 102(3):209–225, 2010.

5. Bahill AT, Clark MR, Stark L. The main sequence, a tool for studying human eye movements. Mathemat Biosci 24(3-4):191–204, 1975.

6. Goossens HH, Van Opstal AJ. Dynamic ensemble coding of saccades in the monkey superior colliculus. J Neurophysiol. 95(4):2326–2341, 2006.

7. Goossens HH, van Opstal AJ. Optimal control of saccades by spatial-temporal activity patterns in the monkey superior colliculus. PLoS Comput Biol. 8(5):e1002508, 2012.

8. Van Opstal AJ, Kasap B. Maps and sensorimotor transformations for eye-head gaze shifts: Role of the midbrain superior colliculus. Prog Brain Res. 249:19–33, 2019.

9. Ottes FP, Van Gisbergen JA, Eggermont JJ. Visuomotor fields of the superior colliculus: a quantitative model. Vision Res 26(6):857–873, 1986.

10. Van Gisbergen JAM, Van Opstal AJ, Tax AAM. Collicular ensemble coding of saccades based on vector summation. Neuroscience 21: 541–555, 1987

11. Kasap B, van Opstal AJ. Modeling auditory-visual evoked eye-head gaze shifts in dynamic multisteps. J Neurophysiol. 119(5):1795–1808, 2018.

12. Quessy S, Freedman EG. Electrical stimulation of rhesus monkey nucleus reticularis gigantocellularis I. Characteristics of evoked head movements. Exp. Brain Res. 156: 342–356, 2004

13. Quessy S, Quinet J, Freedman EG. The locus of motor activity in the Superior Colliculus of the rhesus monkey is unaltered during saccadic adaptation. J. Neurosci. 30: 14235–14244, 2010

14. Walton MMG, Freedman EG: Gaze-shift duration, independent of amplitude, influences the number of spikes in the burst for medium-lead burst neurons in pontine reticular formation. Exp. Brain Res. 214: 225–239, 2011

15. Goodman D, Brette R. Brian: a simulator for spiking neural networks in python. Front Neuroinform. 2:5, 2008.

16. Kasap B, van Opstal AJ. A spiking neural network model of the midbrain superior colliculus that generates saccadic motor commands. Biol Cybern. 111(3-4):249–268, 2017.

17. Alizadeh A, Van Opstal AJ. A spiking neural network model of the Superior Colliculus that is robust to changes in the spatial-temporal input. Sci Rep. 12(1):6916, 2022.

18. Robinson DA. Eye movements evoked by collicular stimulation in the alert monkey. Vision Res 12(11):1795–1808, 1972.

19. Brette R, Gerstner W. Adaptive exponential integrate-and-fire model as an effective description of neuronal activity. J Neurophysiol. 94(5):3637–3642, 2005.

20. Touboul J, Brette R. Dynamics and bifurcations of the adaptive exponential integrate-and-fire model. Biol Cybernet 99(4-5):319–334, 2008.

21. Van Opstal AJ, Hepp K, Suzuki Y, Henn V. Influence of eye position for the collicular role in saccade generation. J Neurophysiol 74: 1593–1610, 1995

22. Scudder CA. A new local feedback model of the saccadic burst generator. J Neurophysiol 59(5):1455–1475, 1988.

23. Robinson DA. Models of the saccadic eye-movement control system. Kybernetik 14, 71–83, 1973

24. Cannon SC, Robinson DA. Loss of the neural integrator of the oculomotor system from brain stem lesions in monkey. J Neurophysiol 57(5): 1383–1409, 1987

25. Laurutis VP, Robinson DA. The vestibulo-ocular reflex during human saccadic eye movements. J Physiol. 373: 209–233, 1986

26. Whittington DA, Lestienne F, Bizzi E. Behavior of pre-oculomotor burst neurons during eye-head coordination. Exp Brain Res 55(2): 215–222, 1984

27. Tomlinson RD, Bahra PS. Combined eye-head gaze shifts in the primate. I. Metrics. J Neurophysiol. 56(6): 1542–1557, 1986

28. Peel TR, Dash S, Lomber SG, Corneil BD. Frontal eye field inactivation alters the readout of superior colliculus activity for saccade generation in a task-dependent manner. J Comput Neurosci. 49, 229–249, 2020

29. Harris CM. Does saccadic undershoot minimize saccadic flight-time? A Monte Carlo study. Vision Res 35(5): 691–701, 1995.

30 Harris CM and Wolpert DM. Signal-dependent noise determines motor planning. Nature 394: 725–726, 1998

31. Tanaka H, Krakauer JW, Qian N. An optimization principle for determining movement duration. J Neurophysiol. 95, 3875–3886, 2006.

32. Van Beers RJ. Saccadic eye movements minimize the consequences of motor noise. PloS one. 3, e2070, 2008

33. Shadmehr R, Mussa-Ivaldi S. Biological learning and control: how the brain builds representations, predicts events, and makes decisions. Chapters 10–12. Mit Press; 2012

34. Sağlam M, Lehnen N, Glasauer S. Optimal control of natural eye-head movements minimizes the impact of noise. J Neurosci 31(45): 16185–16193, 2011.

35. Kardavakis AA, Moschiovakis AK. Optimal control of gaze shifts. J Neuroscience 29 (24): 7723–7730, 2009

36. Kasap B, van Opstal AJ. Dynamic parallelism for synaptic updating in GPU-accelerated spiking neural network simulations. Neurocomp 302:55–65, 2018.

37. Tweed D, Vilis T. Implications of rotational kinematics for the oculomotor system in three dimensions. Journal of Neurophysiology. 58(4): 832–849, 1987

38. John A, Aleluia C, Van Opstal AJ, Bernardino A. Modelling 3D saccade generation by feedforward optimal control. PLoS Comp Biol 17(5): e1008975, 2021

39. Glenn B, Vilis T. Violations of listing’s law after large eye and head gaze shifts. J Neurophysiology, 68(1):309–318, 1992.

40. Tweed D, Haslwanter T, Fetter M. Optimizing gaze control in three dimensions. Science 281 (5381):1363–1365, 1998.

41. Tweed D. Three-dimensional model of the human eye-head saccadic system. J Neurophysiology, 77(2):654–666, 1997

42. Daemi M, Crawford JD. A kinematic model for 3-d head-free gaze-shifts. Front Comput Neurosci, 9:72, 2015

43. Van Opstal AJ, Hepp K, Hess BJ, Straumann D, Henn V. Two-rather than three-dimensional representation of saccades in monkey superior colliculus. Science, 252(5010):1313–1315, 1991

44. Crawford JD, Guitton D. Visual-motor transformations required for accurate and kinematically correct saccades. J Neurophysiology, 78:1447–1467, 1997

